# Genomic prediction of arsenic tolerance and grain yield in rice. Contribution of trait-specific markers and multi environment models

**DOI:** 10.1101/2020.09.28.316356

**Authors:** Nourollah Ahmadi, Tuong-Vi Cao, Julien Frouin, Gareth J. Norton, Adam H. Price

## Abstract

Many rice-growing areas are affected by high concentrations of arsenic (*As*). Rice varieties that prevent *As* uptake and/or accumulation can mitigate *As* threats to human health. Genomic selection is known to facilitate rapid selection of superior genotypes for complex traits. We explored the predictive ability (PA) of genomic prediction with single-environment models, accounting or not for trait-specific markers, multi-environment models, and multi-trait and multi-environment models, using the genotypic (1600 K SNP) and phenotypic (grain arsenic content, grain yield and days to flowering, observed under two irrigation systems over two years) data of the Bengal and Assam *Aus* Panel (BAAP). Under the base-line single environment model, PA of up to 0.707 and 0.654 was obtained for grain yield and grain *As* respectively, the three prediction methods (BL, GBLUP and RKHS) considered performed similarly, and marker selection based on linkage disequilibrium allowed to reduce the number of SNP to 17 K, without negative effect on PA of genomic predictions. Single environment models giving distinct weight to trait-specific markers in the genomic relationship matrix outperformed the base-line models up to 32%. Multi-environment models, accounting for G × E interactions, and multi-trait and multi-environment models outperformed the base-line models by up to 47% and 61%, respectively. Among the multi-trait and multi-environment models, the Bayesian multi-output regressor stacking function obtained the highest PA (0.831 for grain *As*) with much higher efficiency for computing time. These findings pave the way for breeding for *As*-tolerance in the progenies of biparental crosses involving members of the BAAP. It also applies to breeding for other complex traits evaluated under multiple environments.

## Introduction

The high concentration of toxic compounds of arsenic (*As*) in the rice grains, is an important problem in many rice-growing areas (Zavala and Duxbury 2008; Brammer and Ravenscroft 2009; Stroud et al. 2011). One of the effective ways of limiting the potential threats to human health it represents is the development of *As* tolerant rice varieties that limit *As* uptake and /or its translocation to the grains. The *O. sativa* species is endowed with a large genetic diversity for *As* accumulation in the grains when grown in presence of high concentrations of *As* (Dasgupta et al. 2004; Rahman et al. 2007; Norton et al. 2009; Norton et al. 2012; Frouin et al. 2019). Several quantitative trait loci (QTL) controlling the accumulation of *As* in rice grains (G*As*) were mapped using recombinant inbred lines (Norton et al. 2010; Norton et al. 2012b; Kuramata et al. 2013; Zhang et al. 2014). Likewise, a large number of loci associated with G*As* were detected with genome wide association analysis (GWAS) (Norton et al. 2014; Frouin et al. 2019; Norton et al. 2019). Unfortunately, to our knowledge, so far, these findings have not translated into marker-assisted breeding for *As*-tolerance in rice, probably because of the large number of QTLs involved and their rather small individual effect. For instance, Norton et al. (2019) detected 74 QTLs for G*As* with individual effect not exceeding 9.9%.

Using a diversity panel of 228 accessions as training set and 95 elite lines as candidate set, Frouin et al. (2019) analysed the predictive ability (PA) of G*As* QTLs detected by GWAS and the PA of genomic prediction. Only 8% of the QTLs detected by GWAS in the training set could be validated in the candidate set. On the other hand, the PA of genomic prediction for G*As*, across populations, reached 0.496. The authors concluded that genomic selection (GS) is the most effective molecular breeding option for the improvement of *As*-tolerance in rice.

Unlike conventional marker-assisted selection, that evaluates the breeding value using only marker data at loci associated with the mapped QTLs, under GS the breeding value is estimated using all marker data available (Meuwissen et al. 2001; Heffner et al. 2009; Lorenz et al. 2011). A reasonably high predictive ability of the genomic estimate of breeding value (GEBV) was obtained in maize (Bernardo and Yu 2007), wheat (Bassi et al. 2015), barley (Sorrells 2015) and oats (Asoro 2011). In rice, moderate to high predictive ability of GEBV was achieved for a large set of traits, using different types of references and breeding populations (for a review, see Ahmadi et al. 2020). For instance, Ben Hassen et al. (2017) reported the feasibility of accurate GEBV for the progenies of biparental crosses in rice, with a reference population composed of a diversity panel that includes the parental lines of those crosses. Likewise, comparing the performances of single-environment and multi-environment models that accounted for genotype by environment interactions, Ben Hassen et al. (2018) reported the higher predictive ability of the latter models. They concluded that, when applied to data from managed G × E experiments (E being different levels of a given abiotic stress), multi-environment genomic prediction models allow to breed rice simultaneously for productivity and for adaptation to the target abiotic stress. Lastly, it was shown that accounting for trait-specific markers improves the predictive ability of genomic selection in rice (Bhandari et al. 2019).

The present study aims at evaluating the effectiveness of the above-mentioned options of genomic prediction for the improvement of *As*-tolerance in rice. Using the publically available large genotypic data and phenotypic data for the Bengal and Assam *Aus* panel (BAAP) (Norton et al. 2018), we explore the predictive ability of genomic prediction for G*As*, with single-environment models, accounting or not for trait-specific markers reported in Norton et al. (2019), and the performances of multi-environment models, and multi-trait and multi-environment genomic prediction models. As the phenotypic dataset includes grain yield (GY) and days to flowering (DTF), the scope of the study encompasses all complex traits.

## Materials and Methods

### Phenotypic and genotypic

#### Plant material

The plant material was composed of 220 accessions belonging to the *Aus* genetic group of *O. sativa*. They were extracted from the *Aus* diversity panel of 266 accessions developed by Norton et al. (2018). The selection criterion for the 220 accessions was the availability of phenotypic data for G*As* (Supplementary Table 1).

#### Phenotypic data

Phenotypic data were produced in Mymensingh (24°43’10.64” N; 90°25’35.98”E), Bangladesh during the dry seasons of 2013 and 2014, hereafter referred to as Year-1 and Year-2. Each year, two independent trials corresponding to the two water management systems tested, alternate wetting and drying (AWD) and continued flooding (CF), were conducted. The target traits included days to flowering (DTF), grain yield (GY), grain total arsenic content (G*As*) and shoot total arsenic content (S*As*). The latter trait was measured only in 2013.

Details of the phenotyping procedures are provided in Norton et al. (2017b; 2018). Briefly, in each trial, the experimental design was a randomised complete block with four replicate blocks. The individual plot was composed of one 4m-long row. At maturity, grains and shoots of each accession were hand-harvested from the six central hills of each row.

Methods for the measurement of G*As* and S*As* are described in Norton et al. (2007a and 2019). Briefly, for each individual plot, first, a sample of the harvested grains were de-husked and oven dried, and a sample of the shoots were oven dried and powdered. Then, the dried grain and shoots powder were digested using nitric acid and hydrogen peroxide. Finally, total arsenic content was measured by inductively coupled plasma – mass spectroscopy (ICP-MS). Trace element grade reagents were used for all digests, and certified reference material (CRM) (Oriental basma tobacco leaves [INCT-OBTL-5]) and rice flour [NIST 1568b]) were used for quality control; blanks were also included. All samples and standards contained 10 μg L^−1^ indium as the internal standard.

Phenotypic data used for genomic prediction in the present study are, for each accession and each trait, the adjusted means under each treatment (i.e. AWD and CF) in year-1, and year-2 of the two years’ field experiments.

#### Genotypic data and marker selection for genomic prediction

The genotypic data used in this study were extracted from the “Bengal and Assam *Aus* Panel (BAAP)” SNP database, containing 2,053,863 SNPs (Norton et al. 2018). Using Gigwa software (Sempéré et al. 2019), the BAAP database was filtered for minor allele frequency (MAF > 5%), heterozygosity (H < 5%) and absence of missing data. The filtering yielded 1,603,611 SNPs hereafter referred to as 1600k SNPs dataset. The BAAP database is available as a project called “BAAP” at the SNP-Seek database (http://snp-seek.irri.org/).

Using the 1600k SNP dataset, two types of smaller datasets were extracted with the aim of (i) identifying the threshold of marker density below which the accuracy of GEBV declines, and (ii) analysing the effect of accounting for trait-specific markers on the performances of genomic predictions.

Four markers’ density levels, corresponding to four linkage disequilibrium (LD) thresholds (r^2^ ≤ 0.25, r^2^ ≤ 0.50, r^2^ ≤ 0.75, and r^2^ ≤ 1), were considered. The 1600k SNP, corresponding to the threshold of r^2^ ≤ 1, was filtered for the remaining three LD thresholds based on the following procedure. First, the 192 trait-specific SNP (see next paragraph) were discarded. Then, for each chromosome, the complete pairwise r^2^ matrix was computed. Then, single loci or clusters of loci with pairwise LD with other loci below the threshold were identified. All such individuals were kept, and for each cluster, one locus with the highest MAF was randomly selected. This procedure yielded 17,449 SNPs, 11,677 SNPs and 4,255 SNPs for the LD thresholds of r^2^ ≤ 0.75, r^2^ ≤ 0.50 and r^2^ ≤ 0.25, respectively.

The selection of trait specific markers was based on GWAS results reported by Norton et al. (2018) for DTF and GY, and by Norton et al. (2019) for G*As* and S*As*. For DTF, 2-3 SNPs were selected along each of the 24 chromosomic segments with “notable association” detected in one of the two years of field experiment; the total number of DTF-specific SNP was 64. For GY, two SNPs were selected at the two extremities of each of the 32 chromosomic segments with “notable association” detected in at least one of the two years of field experiment, under at least one of the two water management systems. For G*As* and S*As*, 64 SNP were selected out of the 74 “most significant” SNPs (P-value < 0.0001) detected for at least one of the two traits, in at least one of the two years of experiment and at least under one of the two water management systems; the remaining 10 most significant SNP were not present in the 1600k SNP dataset. The list of the 192 trait-specific markers is provided in Supplementary Table 2.

### Methods for genomic prediction

#### Single environment models

Two kernel regression models and a penalised regression model were used in a Bayesian framework, to predict GEBV: the genomic best linear unbiased prediction (GBLUP), the reproducing kernel Hilbert spaces regressions (RKHS), and the Bayesian least absolute angle and selection operator (Bayesian LASSO or BL). Under the GBLUP (VanRaden 2008), the genetic effects are considered as strictly additive and the model is fitted using the genomic relationship matrix (GRM) G = M*M’ (M being the matrix of marker genotypes) as kernel matrix in the Gaussian process implemented. The RKHS model (Gianola and van Kaam 2008) that do not rely on formal hypotheses on the type of genetic effects, was fitted using the squared-Euclidean distance matrix between accessions (derived from matrix of marker genotypes) as kernel matrix; the bandwidth parameter was equal to 0.5. The BL model (Tibshirani 1996, Park and Casella 2008) that hypothesizes most markers do not have any effect, applies a strong shrinkage of estimates of marker effects toward zero for markers with small effects. The prior density of marker effects has a Gaussian distribution and, unlike non-Bayesian LASSO, BL does not impose limitation on the number of non-zero regression coefficients. The model was fitted with a formulation where the prior on the regularization parameter *λ* is expressed as a *β* distribution that allows expressing vague preferences over a wide range of values of *λ*.

#### Prediction model using trait-specific genomic relationship matrix

Trait-specific GRM (G’) were constructed following the method proposed by Zhang et al. (2014). Briefly, it consists in (i) building the standard GRM of Van Raden (2008) (G), with the set of markers that are assumed to have identical small effects not specific to the phenotypic trait considered. (ii) For each trait, building a specific relationship matrix (S), using markers known for their rather large effect on the trait, while specifying the effect w_i_ of each of those markers. (iii) Finally, building for each trait the weighted GRM as:

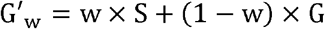

where *w* is the overall weight for trait-specific markers *w* ∈ [0,1]. In the present study, the G matrix was built with the 17,449 SNPs dataset (LD threshold of r^2^ ≤ 0.75). For each trait, four G’_w_ GRM were computed, corresponding to *w* values of 0.25, 0.5, 0.75 and 1, while each of the corresponding 64 trait-specific SNPs were given an equal weight, i.e., 0.25, 0.5, 0.75 or 1. The resulting four weighted trait-specific GRMs (G’_0.25_, G’_0.50_, G’_0.75,_ G’_1_) were used as kernel in the Gaussian process of implementation of GBLUP in a Bayesian framework. The PA of genomic prediction obtained with the four G’_w_ GRMs was compared to the ones obtained with the standard G matrix computed with 17,449 SNP alone (G’_0_ *=* G, i.e. w = 0), and with a second G matrix computed with 17,449 SNP plus the 64 trait-specific SNPs while they were not given specific weight (G’_nw_).

### Multi-environment models

The multi-environment models used to predict the GEBV with data from AWD and CF irrigation systems, are the above-described GBLUP and RKHS endowed with extensions to include the environmental effects. The extended GBLUP model (Lopez-Cruz et al. 2015) separates the effects of each marker into a main effect for all the environments and an effect specific to each environment. The extended RKHS model (Cuevas et al. 2017) hypothesizes correlated performances of accessions in different environments and models the genetic correlation between environments with a matrix of order *m*×*m*, *m* being the number of environments.

#### Multi-trait and multi-environment models

To predict GEBV with data from AWD and CF environments while taking into account the correlation between traits and trait × genotype × environment interaction, we used the Bayesian multi-trait and multi-environment model (BMTME) developed by Montesinos-López et al. (2016), and the Bayesian multi-output regressor stacking (BMORS) function described in Montesinos-López et al. (2019). BMTME can be considered as a Bayesian GBLUP for multiple traits and multiple environments using a linear mixed model that includes the trait × environment (T × E) interaction term as a fixed effect, and the trait × genotype (T × G) interaction and the three-way (T × G × E) interaction term as random effects, assuming independence between environments. The BMORS function does not specify the T × E interactions in a variance-covariance matrix. Instead, using the multi-target regression approach (Spyromitros-Xioufis et al. 2016), it relies on a meta-model. First, for each trait (*t*), a multi-environment GBLUP model is implemented. Then, for each trait a meta-model is implemented to scale the predictions of the first-step model with the predictions of the *t*−1 other first-step models. Each meta-model is implemented with a mixed model where the covariates represent the scaled predictions of each trait obtained with the GBLUP model of first-step analysis. The scaling of each prediction is performed by subtracting its mean and dividing it by its corresponding standard deviation.

#### Computing the predictive ability of genomic prediction

Genomic prediction with single environment models was performed using phenotypic data from each combination of irrigation system (AWD or CF) and year of field experiment (year-1 and year-2), hereafter referred to as AWD1, AWD2, CF1 and CF2. The cross-validation scheme used 80% of the accessions (i.e. 176 accessions) as the training set and the remaining 20% (44 accessions) as the validation set.

The multi-environment models were fitted separately for data from year-1 and year-2 (i.e., AWD1 and CF1on the one hand and AWD2 and CF2 on the other hand). In each case, the PA was computed with two cross-validation schemes. Under the first scheme (CV1), individuals serving as training set (80% of the observations) are all endowed with phenotypic data from the two environments while phenotypic data are lacking for all the individuals included in the validation set (20% of the observations). Under the second scheme (CV2), again 80% of the observations served as training set and the remaining 20% as validation set, however, individuals in both the training set and the validation set are endowed with at least one phenotypic data in at least one environment.

The BMTME model and the BMORS function were fitted first with data from year-1 and year-2 separately, then with data from the two years jointly. Cross validations were performed with the CV2 method, only.

For all models, cross-validation consisted in one hundred replicates of the random partitioning of accessions into training (80%) and validation (20%) sets. For each partition, the Pearson correlation coefficient between the predicted and the observed phenotypes in the validation set was computed. Then, PA of genomic prediction was computed as the average of the 100 replicates of the correlation coefficients. The standard error associated to each PA was also calculated. For the multi-environment models, the correlation was calculated within each environment. The same partitions were used under both GBLUP and RKHS models to compute the PA for DTF, G*As* and GY traits. All models were fitted with 25,000 iterations for the Gibbs sampler and 5,000 burn-ins. In the case of BMTME, which requires very long computing time, the cross-validation was performed with 5,000 iterations and 1,500 burn-ins.

Variance of the computed PA was partitioned into different sources through ANOVA after transformation of the correlation coefficient data into a Z-statistic (Z = 0.5{ln[1 + r]−ln[1 − r]}). The Z-statistic was transformed back to r variable after ANOVA.

Phenotypic data for shoot arsenic content (S*As*) in year-1 were used only under the BMTME model and the BMORS function to predict G*As*.

#### Implementation of the models

The single environment models and the multi-environment were implemented in the R-3.4.2 environment [55] with the R packages *BGLR* 1.0.5 (Pérez and de los Campos 2014). The multi-trait and multi-environment models BMTME and BMORS were implemented in the R-3.6.1 environment with the R package BMTME (Montesinos-López et al. 2019). The analyses were supported by the CIRAD - UMR AGAP HPC Data Centre of the South Green Bioinformatics platform (http://www.southgreen.fr/).

## Results

### Single environment genomic prediction

The PA of genomic predictions in the 144 cross validation experiments involving four levels of LD threshold, (r^2^ ≤ 0.25, r^2^ ≤ 0.50, r^2^ ≤ 0.75, and r^2^ ≤ 1), three prediction methods (BL, GBLUP and RKHS), and three phenotypic traits (DTF, G*As*, GY) observed under two irrigation systems (CF and AWD) over two years, ranged from 0.434 ± 0.109 to 0.708 ± 0.073, with an overall average of 0.563 (Figure 1; Supplementary Table 3). Analysis of the sources of variation of PA revealed that among the factors considered, two had highly significant effects: LD threshold and phenotypic traits. The effects of trait × LD and trait × irrigation-system interactions were also highly significant (Supplementary Table 4). The least significant difference (LSD) test, showed that whatever the trait, the average PA of genomic prediction with LD threshold of r^2^ ≤ 0.25 were systematically lower than PA obtained with the three other LD thresholds. Likewise, whatever the trait, the average PA with r^2^ ≤ 0.75 were either equal or significantly higher than the PA obtained with other LD thresholds, including r^2^ ≤ 1. On the other hand, whatever the LD threshold, average PA for GY were significantly higher than the average PA for TDF and G*As*. The latter systematically had the lowest average PA. The trait × LD interaction was the highest for G*As* and GY. The highest trait × irrigation-system interaction was observed for G*As*.

**Figure 1:**
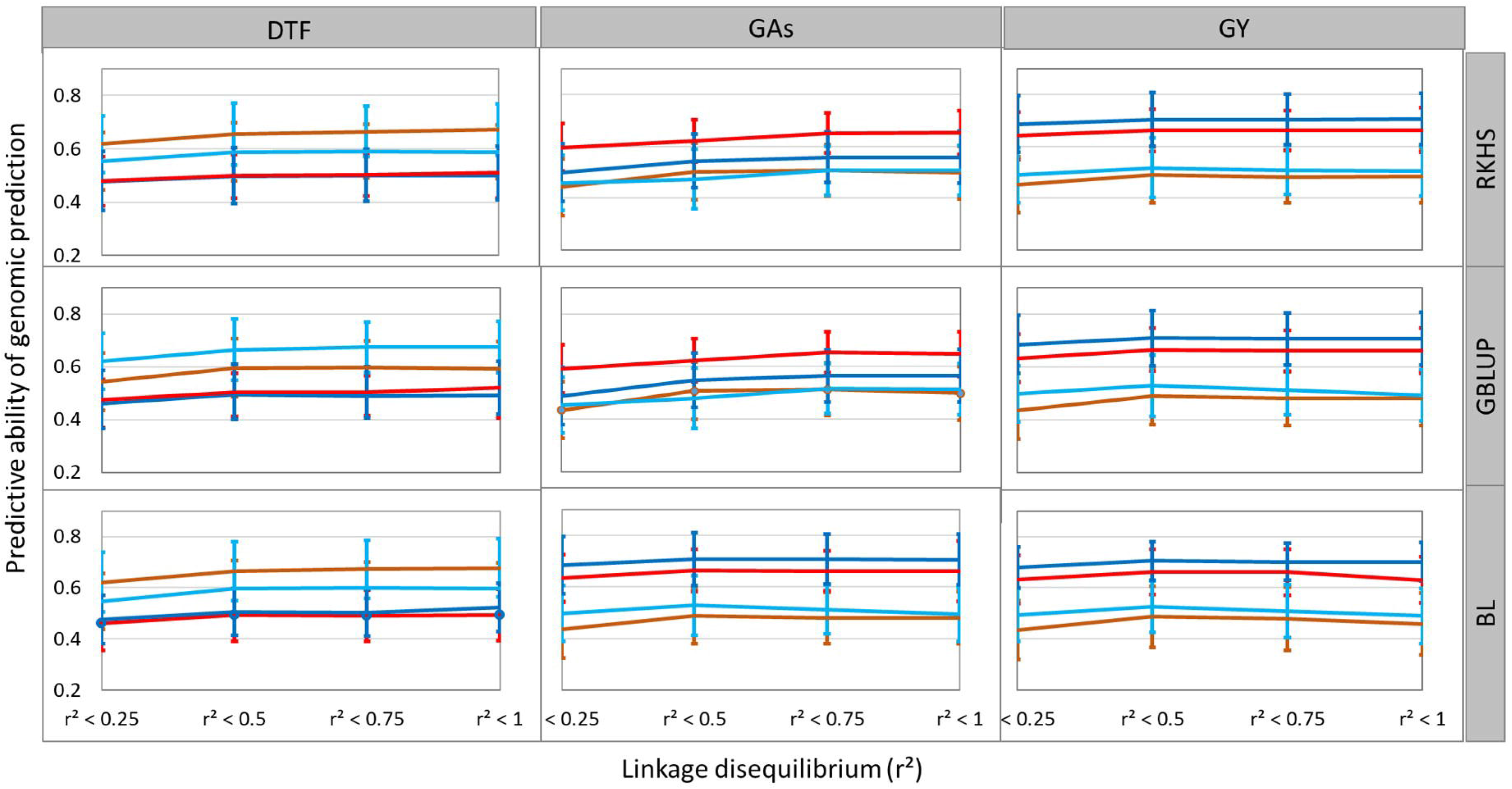
Predictive ability of genomic prediction in cross validation experiments for grain arsenic content (GAs), grain yield (GY) and days to flowering (DTF) observed under alternate watering and drying (AWD) and continued flooding (CF) irrigation systems over two years. Three single-environment statistical methods (BL, GBLUP and RKHS) and four thresholds of linkage disequilibrium between markers (r^2^ ≤ 0.25, r^2^ ≤ 0.50, r^2^ ≤ 0.75 r^2^ ≤ 1) were considered. Light and dark red lines represent AWD year-1 and year-2, respectively. Light and dark blue lines represent CF Year-1 and Year-2, respectively.

Given the systematically high PA obtained with the genotypic dataset selected with LD threshold of r^2^ ≤ 0.75, we decided to use this dataset (17,449 SNPs) for the next steps of our analysis.

### Genomic prediction accounting for trait-specific markers

The 72 cross validation experiments involving six levels of weight of trait-specific markers (G’_0_, G’_nw_, G’_0.25_, G’_0.50_, G’_0.75_, G’_1_) and three phenotypic traits (DTF, G*As*, GY) observed under two environments (AWD and CF) during two years, yielded PA of genomic prediction ranging from 0.480 ± 0.122 to 0.792 ± 0.803. The overall mean PA was 0.671 (Figure 2; Supplementary Table 5). Analysis of the sources of variation of PA revealed that the effects of trait-specific relationship matrix and phenotypic trait were highly significant as well as the effect of interaction between the two factors (Supplementary Table 6). The Tukeys's honest significant difference (HSD) test showed that inclusion of trait specific markers without specific weight did not increase significantly the PA of genomic prediction. The average PA for G’_0_ and G’_nw_ treatments were of 0.573 and 0.584, respectively (Supplementary Table 6). The HSD test also showed that all G’_nw_ GRMs, led systematically to significantly lower PA than GRMs with weighted trait-specific markers (average PA ranging from 0.708 to 0.725). Whatever the environment and the trait, the G’_0.50_ GRM systematically had the highest PA (average PA of 0.725) among the four weighted G’_w_. Likewise, whatever the environment and the trait, the G’_1_ had, almost systematically, the lowest PA (average PA of 0.708) among the four treatments with weighted specific markers. Among the three traits, DTF was the most sensitive to accounting for trait specific markers. Compared to G’_0_ treatment, the average increase of PA for the four GRMs accounting for trait specific markers was 32% for DTF, 19% for G*As* and 28% for GY.

**Figure 2:**
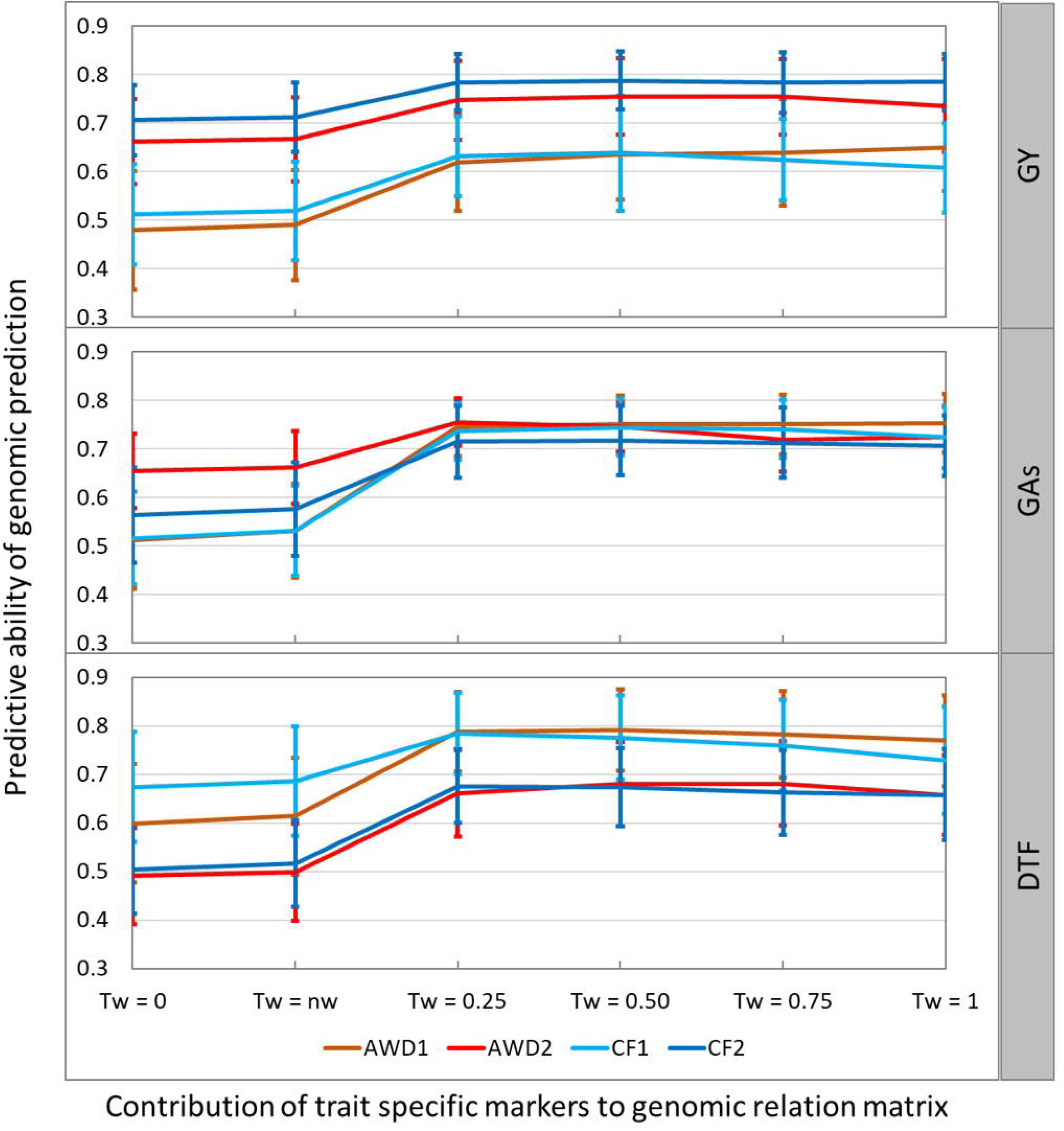
Effects of presence and weight of 64 trait-specific markers on the predictive ability of genomic prediction for grain arsenic content (GAs), grain yield (GY) and day to flowering (DTF) observed in four environments. The three traits are evaluated in four environments: AWD1 and AWD2: Alternate watering and drying Year1 and Year2, respectively; CF1 and CF2: continuous flooding Year-1 and Year-2, respectively. *w* = 0: Establishment of the trait-specific Genomic relationship matrix G’ with 17k SNP; *w* = n*w*: Establishment of Genomic relationship matrix with 17k SNP + 64 trait-specific marker; *w* = 0.25, *w* = 0.50, *w* = 0.75 and *w* = 1: Establishment of Genomic relationship matrix with 17k SNP + 64 trait-specific markers with a weight of 0.25, 0.50, 0.75 and 1, respectively.

### Predictive ability of genomic prediction using multi-environment models

The 72 cross validation experiments involving three levels of models (SE, ME-CV1, and ME-CV2), two prediction methods (GBLUP and RKHS) and three phenotypic traits (DTF, G*As*, GY), observed under two environments (CF and AWD) over two years, yielded PA of genomic prediction ranging from 0.482 ± 0.119 to 0.812 ± 0.050. The overall mean PA was 0.629 (Figure 3; Supplementary Table 7). Analysis of the sources of variation of PA revealed highly significant effects for model and trait factors (Supplementary Table 8). Several interactions of order two, including trait × model, and of order three, including trait × model × method, were also significant. The multi-environment model with the CV2 cross validation strategy significantly outperformed single environment model with an average gain of PA of 0.182 and 0.161 for DTF, under AWD and CF, respectively. The gain of PA was 0.195 and 0.214 for G*As* and 0.164 and 0.150 for GY under AWD and CF, respectively. On the other hand, the PA obtained with ME-CV1 was not significantly higher than the PA obtained with single environment (SE) model. Similar to the trend observed with SE model, the average PA for GY under one of the two ME models was often higher than the PA for DTF and G*As*. The significant effect of trait × model interaction reflected the ability of ME-CV1 to achieve significantly higher PA than SE model for GY, which was not the case for DTF and GY. The trait × model × method interaction reflected, mostly, the fact that under GBLUP method the ME-CV1 achieved the most discriminant PA among the three traits, while under RKHS method the most discriminant PA among the three traits was achieved by ME-CV2 (Supplementary Table 8).

**Figure 3:**
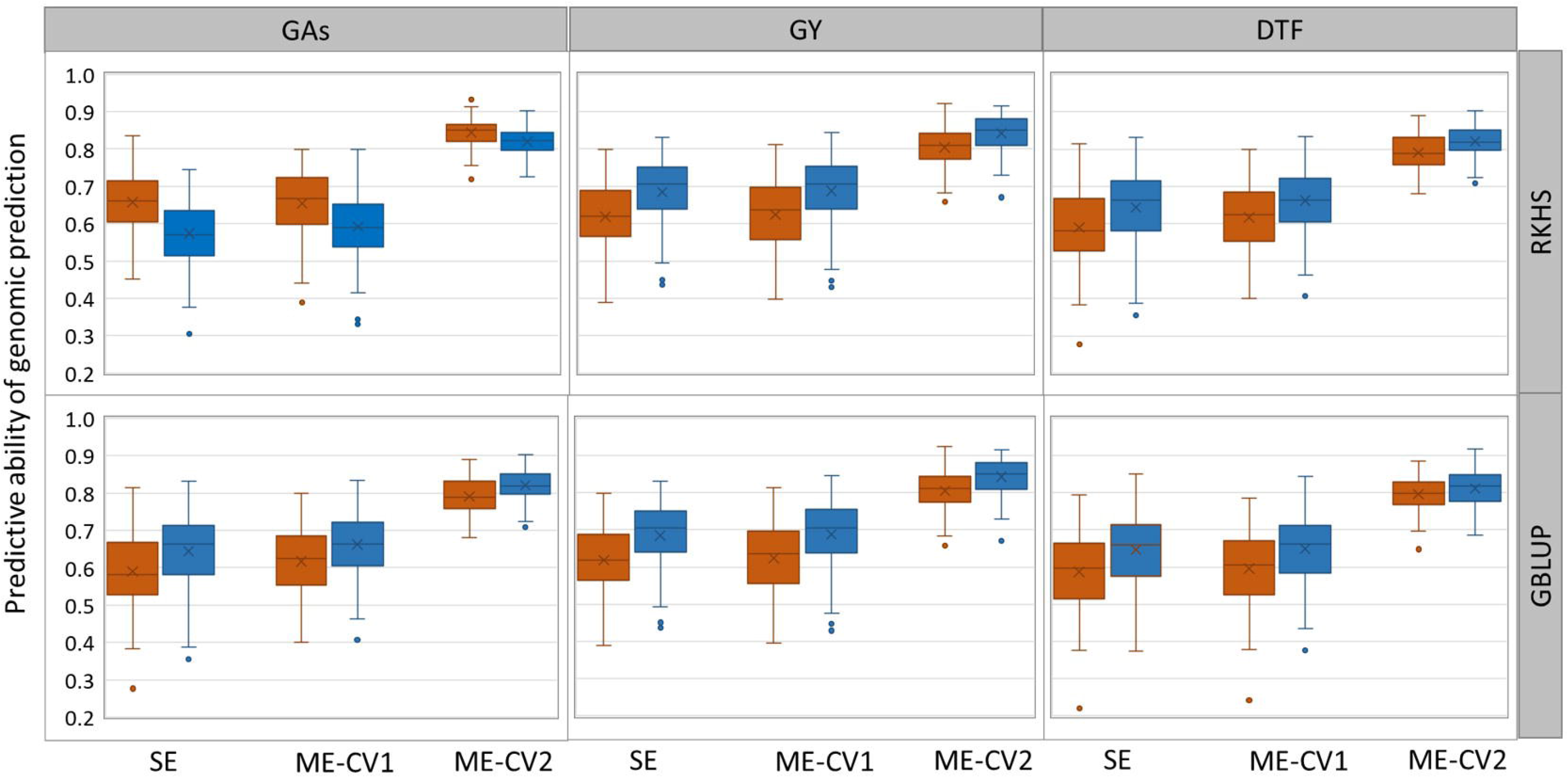
Predictive ability of genomic prediction experiment with single environment (SE), and multi-environment (ME) models obtained with the GBLUP and RKHS statistical methods, for grain arsenic content (GAs), grain yield (GY) and day to flowering (DTF).The ME models are implemented with two cross-validation strategies CV1 and CV2. Two environments are considered: alternate watering and drying (AWD) in orange and continuous flooding (CF) in blue. For each box, the mean (x) and median (horizontal bar) values are represented.

Investigating the possible benefit of a larger number of environments, we implemented the ME-CV2 under GBLUP, while considering each combination of irrigation system and year of experiment as one environment. This provided four environments and allowed predicting traits in each of the four environments (AWD1, AWD2, CF1 or CF2) while accounting for marker effects in the three remaining environments (i.e. predicting AWD1, using information from AWD2, CF1 or CF2, and so on). The effect of a larger number of environments on the PA of genomic prediction was significantly positive (Supplementary Table 9) but modest, always below 10% (Supplementary Table 7). The effects of E × T interaction were also highly significant. For instance, the average PA for G*As* using four environments was +9% under CF and −3% under AWD.

Investigating the possible benefit of accounting for trait-specific marker in a multi-environment model, we compared the PA of ME-CV2 under GBLUP using a genomic relationship matrix computed with the 17k SNP plus 64 trait-specific markers, the latter either with non-specific weight or with a weight of 0.5 (G’_0_ or G’_0.5_). The effect of accounting for trait-specific markers on the PA of genomic prediction was significantly positive (Supplementary Table 10), but very modest, 4% on average and always below 10%. (Supplementary Table 7).

### Predictive ability of genomic prediction using multi-trait & multi-environment models

The multi-trait and multi-environment models were implemented under two options of number of environments: two environments where data from Year-1 and Year2 were analysed separately; four environments, where data from the two years were analysed jointly.

Whatever the number of environments, genetic correlations between traits were very low and negative between DTF and GY, almost nil between DTF and G*As*, very low between G*As* and GY (Table 1). The residual correlation between traits was also low, but could reach 80% of the corresponding genetic correlation between traits. In contrast, correlations between environments were high and always above 0.600 (Table 2). Joint analysis of the two years’ data markedly improved the correlation between the AWD and CF environments (Table 2), but not between traits (Table 1).

**Table 1:**
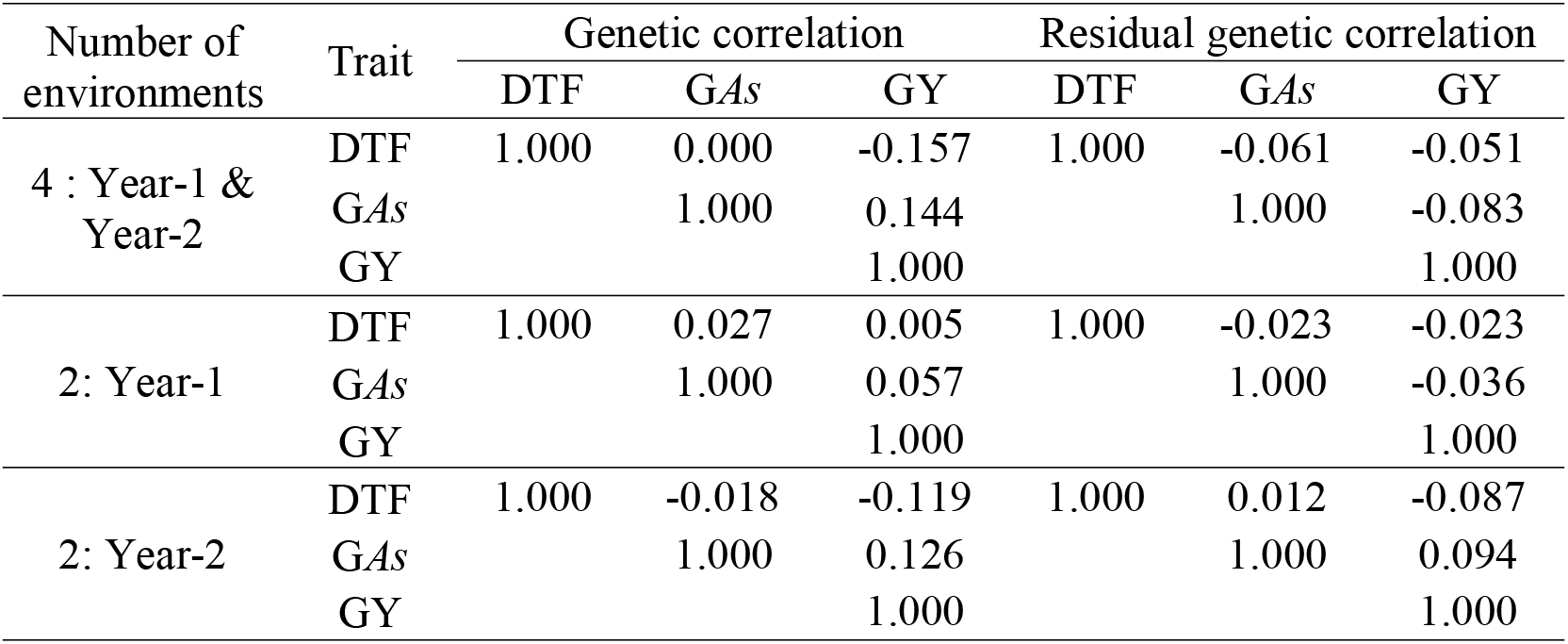
Genetic correlation between days to flowering (DTF), grain arsenic content (G*As*) and grain yield (GY) observed under alternate watering and drying (AWD), and continuous flooding (CF) irrigation systems, over two years.

**Table 2:** Genetic correlation between the alternate wetting and drying (AWD) and the continuous flooding (CF) irrigation systems.

| Number of environments | Irrigation system | Genetic correlation between environments |  |  |  |
| --- | --- | --- | --- | --- | --- |
|  |  | AWD1 | AWD2 | CF1 | CF2 |
| 4 : Year-1 & Year-2 | AWD1 | 1.000 | 0.642 | 0.673 | 0.607 |
|  | AWD2 |  | 1.000 | 0.706 | 0.814 |
|  | CF1 |  |  | 1.000 | 0.639 |
|  | CF2 |  |  |  | 1.000 |
| 2: Year-1 | AWD1 | 1.000 |  | 0.504 |  |
|  | CF1 |  |  | 1.000 |  |
| 2 : Year-2 | AWD2 |  | 1.000 |  | 0.461 |
|  | CF2 |  |  |  | 1.000 |

Results after fitting of the BMTME model for the three traits are summarized in Supplementary Figure 1. Correlation between the observed and the predicted values are high: 0.980 for DTF, 0.985 for G*As* and 0.950 for GY. Results of evaluation of predictive ability of BMTME model are summarized in Figure 4 and Supplementary Table 11. Under the four environments’ option, the PA ranged between 0.259 ± 0.133 and 0.654 ± 0.116 for different combinations of traits and environments. The overall average PA was 0.466. Across the four environments, the largest average PA for DTF was 0.514, for G*As* 0.370 and for GY 0.654. Under the two environments’ option the PA were lower than under the four environments’ option, with an average decrease of 10 %.

**Figure 4:**
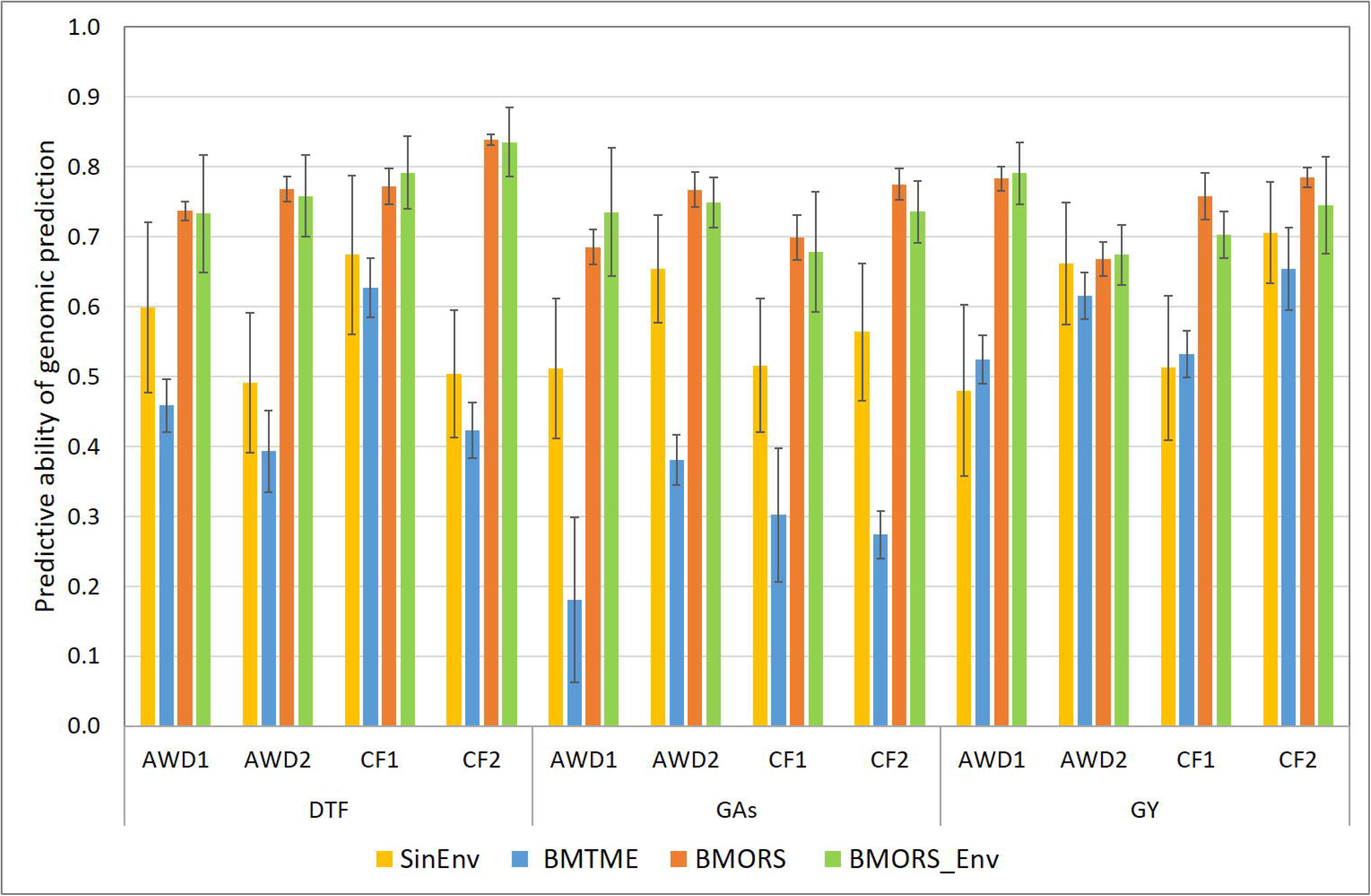
Predictive ability of genomic prediction experiment with three multi-trait and multi-environment prediction models. BMTME: Bayesian multi-trait and multi-environment; MORS: Bayesian multi-output regressor stacking; BMORS_Env: BMORS that allow predicting whole environments using the remaining environments as training. Four environments are considered: alternate watering and drying (AWD1 and AWD2) and continuous flooding (CF1 and CF2). GAs: grain arsenic; GY: grain yield; DTF: days to flowering.

Predictive ability of the BMORS function under the four environments’ option ranged between 0.688 ± 0.074 and 0.839 ± 0.079 for different combinations of traits and environments (Supplementary Table 11). The overall average PA was 0.779. Across the four environments, the largest PA was 0.839 ± 0.079 for DTF, 0.831 ± 0.048 for G*As* and 0.785 ± 0.069 for GY. Under the two environments’ option the PA achieved with the BMORS function were slightly lower than under the four environments’ option, with an average decrease of 4 %. Implementation of BMORS_Env function that allows predicting the traits in each environment, using the remaining environments as training, did not improve the function’s PA but increased the standard deviation associated to each PA (Supplementary Table 11). The overall average PA was 0.744. Across the four environments, the largest PA was 0.835 ± 0.121 for DTF, 0.792 ± 0.169 for G*As* and 0.791 ± 0.165 for GY. Overall, the BMORS function produced significantly higher PA than the BMTME model and the single environment models (Supplementary Table 12).

Taking advantage of phenotypic data available for shoot arsenic content (S*As*) in year-1 of our experiments, we investigated the PA of BMTME model and the BMORS function for G*As*, in the context of the moderate genetic correlation (0.361) between the S*As* and G*As*. The PA of BMTME fitted with S*As* and G*As* data in year-1 was 0.652 ± and 0.659 for G*As* under AWD1 and CF1, respectively. These PA were significantly higher than the largest PA of 0.370 obtained for G*As*, when the model was fitted with G*As* plus DTF and GY data. The PA of BMORS for G*As*, was 0.752 under AWD1, and 0.765 under CF1.

## Discussion

Breeding *As*-tolerant rice varieties capable of limiting *As* uptake and/or its translocation to the grains can mitigate the threat to human health associated with high soil or irrigation water *As* concentration in many rice growing areas. The objective of this study was to take advantage of the availability of a robust phenotypic dataset on the grain arsenic content of the *Aus* rice genetic group, associated with a very large genotypic dataset, to analyse the performances of different genomic prediction approaches for the reduction of grain arsenic.

### Marker density and use of trait-specific markers

Analysis of the effect of four levels of marker density, on the PA of three single-environment genomic prediction models, confirmed the superfluity of very dense genotyping when the LD extent is large. Indeed, simulation studies (Habier et al. 2009; Zhong et al. 2009) and empirical studies in various plant species including rice (Ben Hassen et al. 2017; Bhandari et al. 2019) have shown very little, if any, effect of marker densities, as long as the distance between adjacent markers remains below the extent of LD. In the case of our *Aus* population the lowest and highest per-chromosome LD decay were 157 Kb and 499 Kb, respectively (Norton et al. 2018). Thus, given the size of the rice genome (430 Mb), the theoretical minimum number of markers needed would be 3 K evenly distributed SNPs. However, the 4 K SNPs dataset (LD threshold of r^2^ ≤ 0.25) led, systematically, to lower PA than the ones with higher numbers of SNPs. This is probably due to the uneven distribution of the SNPs, and to local variation of LD along each chromosome. The 17 K dataset (LD threshold of r^2^ ≤ 0.75) provided PA equal to the one observed with the 1600 K dataset. Its SNP density, compensated for the unevenness of their distribution and for the possible low local LD in each chromosome.

Whatever the size of the genotypic dataset and the phenotypic trait considered, the PA of genomic prediction obtained with the three prediction methods (BL, GBLUP and RKHS) were not significantly different. Given the fact that several QTL of rather large effect (up to 5% for DTF, 10% for G*As* and 25% for GY) were detected in the BAAP population (Norton et al. 2018; 2019), one would expect the BL method to perform better than GBLUP and RKHS. This is probably due to the strong linkage of each QTL of rather large effect with a large number of SNPs, taking off the shrinkage edge of BL method. Indeed, the total number of SNP associated with DTF, G*As* and GY was of 3,515 SNPs, 2,720 SNPs and nearly 1,500, respectively (Norton et al. 2018; 2019).

Exploitation of the results of recent GWAS (Norton et al. 2018; 2019), via genomic prediction using trait-specific GRM under GBLUP method, was awarded with PA gains ranging from 19 to 32%. Similar gains of PA were reported in dairy cattle for milk yield (Zhang et al. 2014) and in rice for traits related to drought tolerance (Bhandari et al. 2019). Using GRM built with markers identified through GWAS in Holstein-Friesian bulls, Veerkamp et al. (2016) reported no increase in the accuracy of genomic predictions compared to GRM built with a large set of markers including or not trait-specific SNPs. They attributed the finding to the small size of the effective population and the associated long extent of LD. On the other hand, in their study, the share of total phenotypic variance explained by the trait-specific markers did not exceed 19% and they did not attribute a specific weight to those markers. In our study, the share of total variance explained by the trait-specific markers was much higher. For instance, for G*As*, the three most important QTLs (represented by 8 SNPs out of the 64 G*As*-specific SNPs) each explained 10 to 18% of the total variations. It is also noteworthy that the simple inclusion of the trait-specific SNPs in the genotypic dataset does not improve the PA of genomic predictions. Attribution of a distinct weight is necessary. We empirically opted for four levels of trait-specific weight and, at each level, attributed the same weight to each of the 64 trait-specific SNPs, regardless of the effect of the underlying QTL. The highest PA, for the three traits considered, was achieved with the trait-specific weight of 0.50. Attribution of weight to trait-specific markers can probably be fine-tuned, by considering the effect of each QTL.

### Accounting for G × E interactions and for correlation between traits

The multi-environment models, using joint data from AWD and CF irrigation systems, with the CV2 cross validation strategy provided gains of PA of genomic prediction ranging between 18% and 47% according to traits and environments. Cuevas et al. (2016; 2017) reported similar higher PA of multi-environment models, compared to single-environment models, using data from multi-local and multi-year trials of maize and wheat. In rice, Ben Hassen et al. (2017), using DTF and GY data from two managed environments (AWD and CF), reported PA gains of up to 30% by using multi-environment models compared to their single-environment counterparts. Likewise, Bhandari et al. (2019) using rice DTF and GY data from three managed drought experiments, reported PA gains of up to 32% with the RKHS multi-environment models. In the present study, the two multi-environment models (GBLUP and RKHS) provided similar substantial gains of PA, despite the fact that under the M×E model implemented with GBLUP, the environmental covariance is forced to be positive and constant across pairs of environments. This is probably because, in our study, correlations between the AWD and CF environments are positive for the three traits, over the two years of the experiment. For instance, in the case of G*As*, the correlation between AWD and CF was 0.709 and 0.754 in year-1 and year-2, respectively. When the correlation between environments is negligible or negative, the G × E model cannot explain the interaction because the sample correlation is estimated through variance components that are, by definition, all positive or zero. Therefore, the G × E model’s prediction accuracy is similar to that of the single-environment model because the variance of the marker main effects tends to be small and positive. Then, negative correlations between environments are not used to improve the predictions.

Predictive ability of the multi-trait and multi-environment model (BMTME), that includes a three-way interaction term (T × G × E), and accounts for correlation between traits, was almost systematically lower than the PA of its single-trait and single-environment counterpart. It was also lower than the PA of the multi-environment models GBLUP and RKHS. This is most probably due to the very low correlation (always below 0.150) between the three traits considered, and the fact that our data did not comply with the hypothesis of unstructured variance-covariance matrix for both genetic and residual covariance matrix between traits. The significantly higher PA obtained when the model was fitted with the more tightly correlated traits G*As* and S*As*, underline the determining effect of the level of correlation between traits. Similar results were reported by the developers of the BMTME model (Montesinos-López et al. 2016), who obtained the highest PA when correlation between traits was above 0.5, in both real and simulated data.

Whatever the trait and the environment, the BMORS function outperformed significantly its BMTME and its single-trait and single-environment models’ counterparts implemented with GBLUP method. This is probably because the first stage models of BMORS are simply univariate multi-environment GBLUP that do not use information on correlation between traits, and the second stage models, though based on hypotheses of correlation between traits, does not directly involve a variance-covariance matrix. Instead, it involves an additional training stage where, for each trait, a meta-model is trained with an expanded training set, composed of the input vectors of the first stage, augmented by the estimates of the values of their target variables for all traits, in the first stage models (Spyromitros-Xioufis et al. 2016).

### Implication for breeding for complex traits

In this study, we showed (i) contrary to widespread idea, more markers is not always better for GS, the number should be adjusted to LD within the population. (ii) Combining the QTL models of infinitesimal effects and large effects (i.e. accounting for trait-specific markers) provides better predictions than each of the two models implemented separately. (iii) Prediction models that account for G × E interactions and/or for correlation between traits achieve higher PA than the conventional single-trait and single-environment models. The three findings applied to both complex traits such as GY and G*As* with moderate heritability, and to DTF, a trait of less complex genetic architecture and high heritability. These findings were based on cross-validation experiments within a diversity panel. Ben Hassen et al. (2017) have reported that reasonably high PA of GEBV could be achieved in rice for the progenies of biparental crosses with a reference population composed of a diversity panel that includes the parental lines of those crosses. Thus, our findings would apply, at least, to progeny of crosses between accessions of the BAAP panel for the improvement of grain yield and lowering grain arsenic in the rice *Aus* group. Furthermore, it applies to all crop-breeding programs that have multi-trait and multi-environment targets.

## Supplementary Material

**Supplementary Table 1:** Phenotypic data of 220 accessions of the Bengal and Assam *Aus* diversity panel used in present study.

**Supplementary Table 2:** List of SNPs associated with the most significant QTLs for days to flowering (DTF), grain arsenic content (GAs) and grain yield (GY) as identified by genome-wide association analyses reported by Norton et al. (2018; 2019).

**Supplementary Table 3**: Predictive ability (PA) of genomic prediction with single environment models BL, GBLUP and RKHS.

**Supplementary Table 4:** ANOVA on predictive ability (PA) of genomic prediction with single environment models.

**Supplementary Table 5:** Predictive ability (PA) of genomic prediction models accounting for trait-specific markers.

**Supplementary Table 6:** ANOVA on predictive ability (PA) of genomic prediction with different weight (T_w_) given to trait specific markers.

**Supplementary Table7**: Predictive ability (PA) of single-environment (SE) and multi-environment (ME) prediction models

**Supplementary table 8**: ANOVA on predictive ability (PA) of genomic prediction with single environment (SE) and multi-environment (ME) models

**Supplementary table 9**: ANOVA on predictive ability (PA) of genomic prediction of multi-environment models using data from two or four environments

**Supplementary table 10**: ANOVA on predictive ability (PA) of genomic prediction with multi-environment models and trait specific markers

**Supplementary table 11**: Predictive ability (PA) of genomic prediction with multi-trait and multi-environment models BMTME and BMORS.

**Supplementary table 12**: ANOVA on predictive ability (PA) of genomic prediction with multi-trait & multi-environment models BMTME and BMORS.

**Supplementary Figure 1:** Graphic representation of the predicted values of three phenotypic traits by the Bayesian multi-trait and multi-environment model (BMTME) against their observed values. DTF: days to flowering; GAs: grain arsenic; GY: grain yield.

