## Supplementary figures and images for "Genomic prediction of arsenic tolerance and grain yield in rice. Contribution of trait-specific markers and multi environment models"

### Supplementary Figure 1.jpg

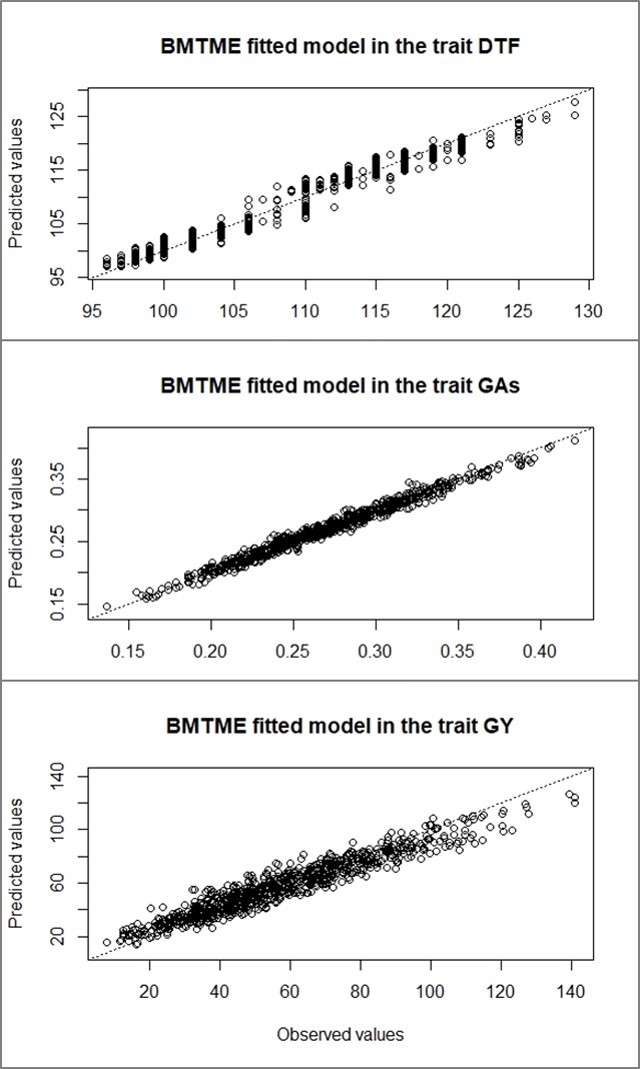
